# A pivotal role for Wnt antagonists in constraining Wnt activity to promote digit joint specification

**DOI:** 10.1101/2025.07.17.665381

**Authors:** Bau-Lin Huang, Sean Davis, Eiki Koyama, Maurizio Pacifici, Susan Mackem

## Abstract

Bmps and Wnts play opposing roles in several contexts during chondrogenesis and joint formation. Using genetic and genomic approaches, we found instead that canonical Wnts can cooperate with Bmps to enhance pSmad1/5 activity and initiate chondrogenic commitment in digit tip progenitors. 5’Hoxd^Δ/Δ^ mutant digits are characterized by elevated Bmp and pSmad1/5 activity and subsequent joint loss. We show that expressing stabilized βcatenin (βcatCA) in interdigit mesenchyme rescues 5’Hoxd^Δ/Δ^ digit joint loss non-autonomously, by inducing secreted Wnt antagonists and normalizing digit tip pSmad1/5 levels. Indeed, genetic removal of *Dkk2* in 5’Hoxd^Δ/Δ^ prevented joint rescue by βcatCA. Furthermore, elevating Wnt activity with Gsk3β antagonists in limb bud culture stabilized pSmad1/5 levels and enhanced Bmp activity. Elevated pSmad1/5, as seen in 5’Hoxd^Δ/Δ^, accelerates chondrogenic commitment, impeding a switch of phalanx forming region (PFR) cells in the digit tip to interzone (joint progenitor) fate. We propose that, before progenitors transit into PFR, Wnt antagonists cooperate with Fgfs to prevent precocious pSmad1/5 accumulation by stabilizing Gsk3β to promote Smad-linker phosphorylation and Smad1/5 degradation. Consequently, Wnt antagonists play a key role in modulating the pace of initial commitment of digit progenitors to chondrogenesis, together with Fgfs, and maintain mesenchymal plasticity to balance digit phalanx and joint formation.

## Introduction

Joints are critical for skeletal mobility and functional limb adaptations, particularly in the hand/foot (autopod) region. The number of phalanges and joints serve as a hallmark of digit identity and vary over a wide range in vertebrates through evolutionary adaption for different functions, such as flying, swimming, running^1^. Understanding the steps regulating phalanx and joint formation offers insights into how early limb patterning is linked to later events in skeletal morphogenesis at the molecular level, as well as illuminating the developmental basis for adaptive evolution. Manipulation of signaling activities of several major pathways, (e.g., Shh, Bmps, Fgfs and Wnts), can alter the position and number of joints and phalanges in the developing digits ^2–5^. Yet how these pathways coordinate *in vivo* to determine digit morphology and joint formation is still poorly understood.

Bmp-, Wnt- and Fgf- are major pathways active at digit-forming stages. The interplay of Fgfs and Wnts maintain mesenchymal progenitors in an undifferentiated and proliferative state that is poised to assume a chondrogenic fate. However, exposure of the mesenchymal progenitors to Wnts in the absence of Fgfs results in stable epigenic changes that suppress *Sox9* activation and chondrogenic fate^6,7^. Bmps are required to promote compaction of limb mesenchymal cells prior to Sox9 expression and the formation of discrete chondrogenic condensations ^8,9^.

Both interzone (joint precursor) cells and chondrocytes are derived from common mesenchymal Sox9+ pre-chondrogenic cells ^10^. However, in contrast to long bones where joints arise following cartilage condensation, digit joint progenitors are specified coordinately with phalangeal precursors at the digit tip in the PFR (phalanx forming region) in response to modulated interdigit Bmp signaling, and to local Noggin. This results in the emergence of sequential, repeating phalanx-interzone anlage in the digit tip ^3,11–13^. Interzone specification entails a loss of *Sox9* RNA and the initiation of *Gdf5* expression^10,14–16^, which serves as the earliest marker of interzone fate. Analysis of mouse mutants, as well as manipulation of signaling in chick, has identified both Wnt ^4,17^ and Bmp ^2,18,19^ pathways as playing pivotal roles in directing this process. Although Wnt/βCatenin activation induces ectopic *Gdf5* expression, *βCatenin* removal in early mouse limb bud does not alter the onset of interzone *Gdf5* expression^20^. In both Noggin and in 5’Hoxd^Δ/Δ^ mutants, *Gdf5* expression never appears in the presumptive joint region of cartilage^2,12^, suggesting that earlier events upstream of Wnt activation may direct initial interzone specification.

To test whether Wnt/βCatenin can initiate interzone specification in developing digits, we selectively expressed a constitutively-active, stabilized βCatenin (βCatCA) ^21^ in 5’Hoxd^Δ/Δ^ cartilage or interdigit tissue. Unexpectedly, βCatCA acted non-autonomously to reduce Bmp response in 5’Hoxd^Δ/Δ^ distal digit tips and restored joint formation by inducing Wnt antagonists, including *Dkk2*. Conversely, interdigit Wnt3a mis-expression resulted in excessive Bmp response and joint loss. In contrast to previous work supporting opposing roles of Bmps and Wnts in joint formation, we show that Wnts cooperate with Bmps by suppressing Gsk3β and stabilizing pSmad1/5, thereby enhancing Bmp activity and promoting the formation and compaction of pre-chondrogenic aggregates from digit mesenchymal progenitor cells to contribute to the PFR. The sustained high level of Bmp activity acts to accelerate transition from compacted PFR to chondrogenic commitment, impairing a switch to interzone fate and consequently joint formation. In this context Wnt antagonists, by stabilizing Gsk3β, serve to slow the rate of compaction/condensation and thereby the pace of chondrogenic determination, enabling a fate switch to occur. Therefore, we conclude that, at an early stage, Wnt antagonists play a pivotal role in constraining Wnt activity and maintaining the plasticity of forming PFR cells to adopt either a chondrogenic or interzone fate.

## Results

### Selective interdigital β-Catenin activation restores joint formation in 5’Hoxd^Δ/Δ^ digits

*5’Hoxd* (*Hoxd11, d12, d13*) gene deletion results in brachydactyly (all bi-phalangeal digits) and compromised metacarpal-phalangeal joint formation in forelimb, particularly in digits 3 and 4 (100% frequency) and with lower penetrance in digits 2 and 5 (15-30% frequency) ^12,22,23^(Figure 1c). We previously showed that reducing Bmp activity in 5’Hoxd^Δ/Δ^ autopod restores phalanx and digit joint formation^12^. Unexpectedly, we found that expression of a stabilized, conditional βCatenin allele (hereafter referred to as βCatCA) in digit chondroprogenitors (using Sox9CreER, see Figure 1f), did not rescue joint formation in 5’Hoxd^Δ/Δ^ digits ^12^, contrary to expectations based on prior work showing that canonical Wnt activity promotes joint formation in long bones by antagonizing Sox9 activity and reversing chondrocyte differentiation^17,24,25^. This led us to ask whether canonical Wnt signaling plays any role in digit joint formation analogous to that in long bone joint induction. To confirm and extend our previous results, we compared the effects of expressing conditional βCatCA in selected 5’Hoxd^Δ/Δ^ limb bud regions using different Cre lines to activate βCatCA expression at the time of digit phalanx/interzone specification (tamoxifen at E11.25).

**Figure 1.**
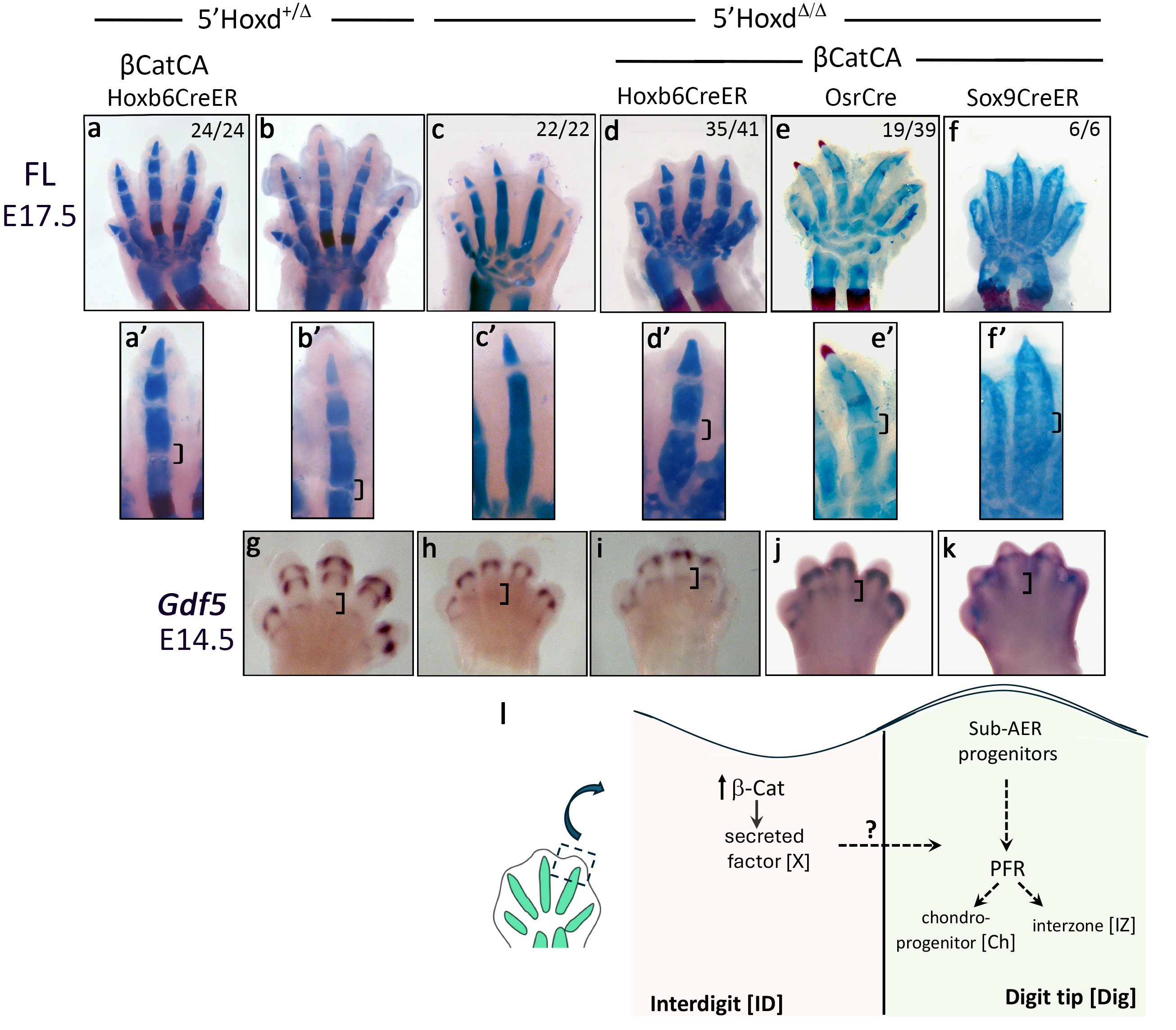
Digit joint formation restored in 5’Hoxd^Δ/Δ^ by βCatCA activation with Hoxb6CreER or OsrCre. **(a-f)** E17.5 limb skeletons show digit morphology with or without indicated Cre- or CreER-activated βCatCA in 5’Hoxd^Δ/Δ^ and control siblings, **(a’-f’)** Images below show joint formation in enlarged digit 3 for each genotype. Bracket indicates the presumptive area of metacarpal-phalangeal joint. The numbers in the images indicate the frequency of observed phenotype in limbs analyzed for each genotype, **(g-k)** in situ hybridization shows Gdf5 expression in interzones. Bracket indicates presumptive area of metacarpal-phalangeal interzone. (I) Schematic of possible non-cell autonomous action of interdigit ID)-activated βCatCA on digit tip (Dig) progenitors, affecting some step in commitment of progenitors to chondrogenic (Ch) or interzone/joint (IZ) fate. See text for details. Tamoxifen injectied at Ell.25 for Hoxb6CreER and Sox9CreER.

We first compared Sox9CreER to Hoxb6CreER, which is expressed in a largely complementary fashion (throughout undifferentiated limb mesenchyme, but shutting off as cells undergo chondrogenic differentiation)^26,27^. As previously reported, chondrogenesis was severely impaired in 5’Hoxd^Δ/Δ^ digits by Sox9CreER-activated βCatCA, but there was no evidence for restoration of interzone/joint formation, either morphologically or by *Gdf5* expression (n=6/6, Figure 1f, 1k compared to 1c, 1h). In contrast, activation of βCatCA by Hoxb6CreER did not perturb gross digit skeletal morphology in control embryos, except for loss of one phalanx in digit 5 (n=24/24, Figure 1a compared to 1b). But compared to cre-negative 5’Hoxd^Δ/Δ^ siblings, metacarpal-phalangeal joint formation was restored in digits 3 and 4 of 5’Hoxd^Δ/Δ^;βCatCA embryos, both at the level of interzone *Gdf5* expression and subsequent skeletal morphology, albeit with mild chondrogenic defects (n=35/41, Figures 1d, 1i, and S1a). However, phalanx restoration (to triphalangeal digits) was never seen (n=35/35). These results suggest that βCatenin-activation in interdigits acts indirectly to affect phalanx/interzone specification in digit progenitors. Inclusion of a Rosa-tdTomato reporter confirmed that Hoxb6CreER activity at the stages used to activate βCatCA is limited predominantly to interdigit mesenchyme with scarce recombinant descendant cells subsequently present in mature cartilage at late stages (E14.5, E16.5) and no contribution to restored joint regions (Figure S2a). To further confirm that rescue was not a result of scarce βCatCA-expressing chondrogenic cells contributing to interzone directly, we used a highly restricted, interdigit-specific OsrCre transgene ^28^ to activate βCatCA expression solely in interdigits, albeit in a domain limited to the digit 3 region. 5’Hoxd^Δ/Δ^ joint formation was also restored by selective interdigit βCatCA expression with OsrCre (n=19/39, Figure 1e, 1j). Owing to the late onset of OsrCre expression in different interdigits (beginning at E12.5 anteriorly from d1 towards d3 and extending to d5 by E13.5^28^, see Figure S2b), OsrCre-activated βCatCA and joint restoration were limited to digit 3, and occurred with lower frequency compared to Hoxb6CreER-driven activation of βCatCA. Lineage analysis with Rosa-tdTomato indicated that OsrCre-activated βCatCA-descendent cells in 5’Hoxd^Δ/Δ^; βCatCA limbs remained completely restricted to interdigit tissue even at very late skeletal stages (E16.5) and no labeled cells were observed in cartilage or interzone/joint regions (Figure S2c). Together, these results indicate that βCatCA acts indirectly from interdigit tissue to promote interzone/joint formation non-cell autonomously (Figure 1l).

Joint formation is a multi-step process, beginning with induction of interzone *Gdf5* expression, and eventually leading to cavitated, mature joints. To confirm that the 5’Hoxd^Δ/Δ^;βCatCA rescue progresses to mature joint formation, we examined the late-stage expression of both *Gdf5* and the articular cartilage-specific factor, *Proteoglycan 4* (*Prg4*), in 5’Hoxd^Δ/Δ^ joints restored by OsrCre-activated βCatCA. In sibling controls, *Gdf5* was maintained in the interzone region from E14.5 on and *Prg4* expression became apparent in the articular cartilage of newborns (P0), but both were absent in 5’Hoxd^Δ/Δ^ digits (Figure S1a,b). In contrast, in 5’Hoxd^Δ/Δ^;βCatCA rescued embryos at E14.5 and P0, respectively, interzone *Gdf5* and articular *Prg4* expression were both restored and joint cavitation was complete at P0 (Figures 1i,j and S1a,b), indicating that the rescued joints persist and mature.

### Rescue of joint formation by interdigit βCatCA expression reduces excess Bmp activity present in the 5’Hoxd^Δ/Δ^ PFR

Previous work in chick has implicated interdigit signals as key regulators that differentially instruct digit identity (final phalanx/joint number) of the adjacent progenitors in digit tip that form the PFR ^3,29^. We previously showed that digit tip PFR cells can adopt either a chondrogenic or joint fate depending on their level of Bmp response and that elevated Bmp activity in the *5’Hoxd^Δ/Δ^* PFR is responsible for inhibiting interzone formation ^12^. To determine if βCatCA acts indirectly to restore *5’Hoxd^Δ/Δ^* interzone formation by modulating Bmp response in the PFR, we analyzed pSmad1/5 levels in distal digit tips in the βCatCA-rescued embryos, compared to 5’Hoxd^Δ/Δ^. In 5’Hoxd^Δ/Δ^ embryos, pSmad1/5 staining was elevated compared to sibling controls, particularly in the sub-AER mesenchyme to distal tip PFR (Figure 2b compared to 2a, brackets), but was reduced in the 5’Hoxd^Δ/Δ^;βCatCA embryos (Figure 2c, bracket). To confirm that interdigit βCatCA restores joint formation by altering the level of Bmp response, we increased relative Bmp levels genetically by reducing the gene dosage of the Bmp antagonist, *Noggin*. Removing one *Noggin* allele (*Nog*^+/−^) in 5’Hoxd^Δ/Δ^;βCatCA-rescued embryos increased pSmad 1/5 level in the PFR and sub-AER regions, as expected (Figure 2d, bracket), and also completely prevented the restoration of digit joint formation (compare digit 3,4 in Figure 2g-arrows, n=13/14 with 2h, n=10/10), indicating that βCatCA restores digit joint formation by modulating the digit tip Bmp response level (Figure 2i).

**Figure 2.**
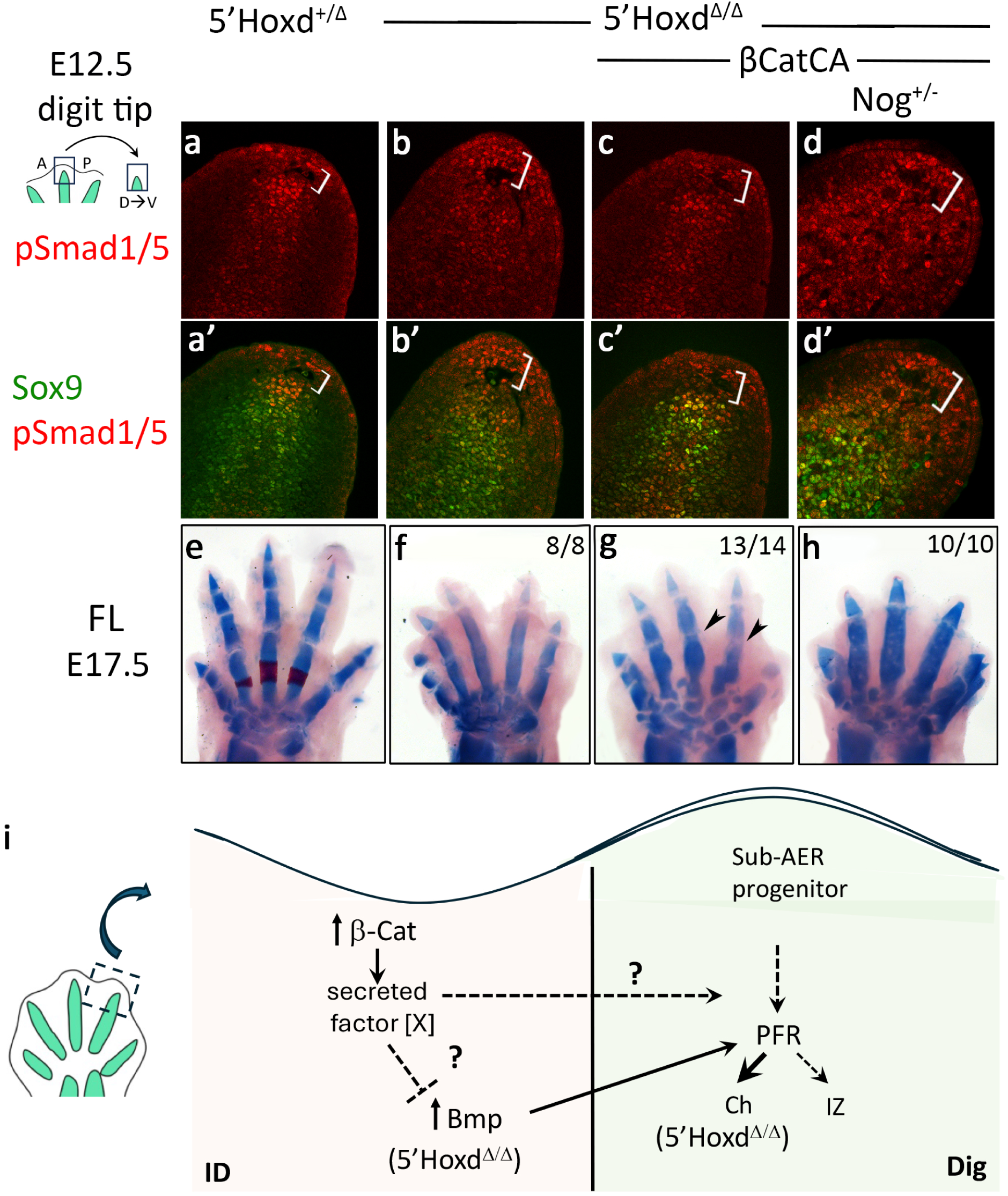
Bmp activity in the digit tip is reduced after βCatCA activation in 5’Hoxd^Δ/Δ^. **(a-d)** pSmadl/5 staining of E12.5 distal digit tip sections (dorsoventral, as shown in schematic) in βCatCA-rescued 5’Hoxd^Δ/Δ^ and effect of reducing Noggin gene dosage. Bracket indicates sub-AER mesenchymal region. **(e-h)** E17.5 skeletal stain showing changes in digit joint formation. Arrowheads indicate digit joints rescued by interdigit βCatCA activation in 5’Hoxd^Δ/Δ^ and loss of rescue following Noggin dosage reduction (compare panels g and h). Numbers in panels indicate frequency of phenotype shown in limbs analyzed. **(i)** schematic summary of possible effects of interdigit (ID)-activated βCatCA on Bmp activity and pace of chondrogenic (Ch) vs interzone (IZ) commitment in digit tips (Dig) of the 5’Hoxd^Δ/Δ^ mutant, which has excess Bmp-pSmad activity that accelerates chondrogenesis and impairs interzone fate. See text for details.

### Interdigit transcriptome analysis identifies Wnt antagonists as candidate βCatCA targets acting non-cell autonomously to restore 5’Hoxd^Δ/Δ^ digit joints

The cell non-autonomous effect of interdigit βCatCA suggested that βCatCA may induce secreted factors acting on PFR cells during joint specification. This prompted us to screen for βCatCA−induced target candidates for restoring 5’Hoxd^Δ/Δ^ digit joint formation using RNAseq analysis of FACS-sorted interdigit cells to compare differential expression (DE) between 5’Hoxd^+/β^ control, 5’Hoxd^Δ/Δ^ and 5’Hoxd^Δ/Δ^;βCatCA sibling embryos. Because of its high rescue efficiency (∼100%), βCatCA expression was activated by Hoxb6CreER, together with Rosa-tdTomato for FACS isolation of interdigit cells (Figure 3a). Principal component analysis showed biological replicates for each genotype clustered separately from other genotypes (Figure 3b). Notably, the 5’Hoxd^Δ/Δ^ and 5’Hoxd^Δ/Δ^;βCatCA replicates clustered more closely with each other than with 5’Hoxd^+/β^ controls, suggesting that interdigit βCatCA altered expression of a limited number of genes in the 5’Hoxd^Δ/Δ^ mutant background, consistent with the DE analysis (Tables S1, S2 and summarized in Figure 3c,d).

**Figure 3.**
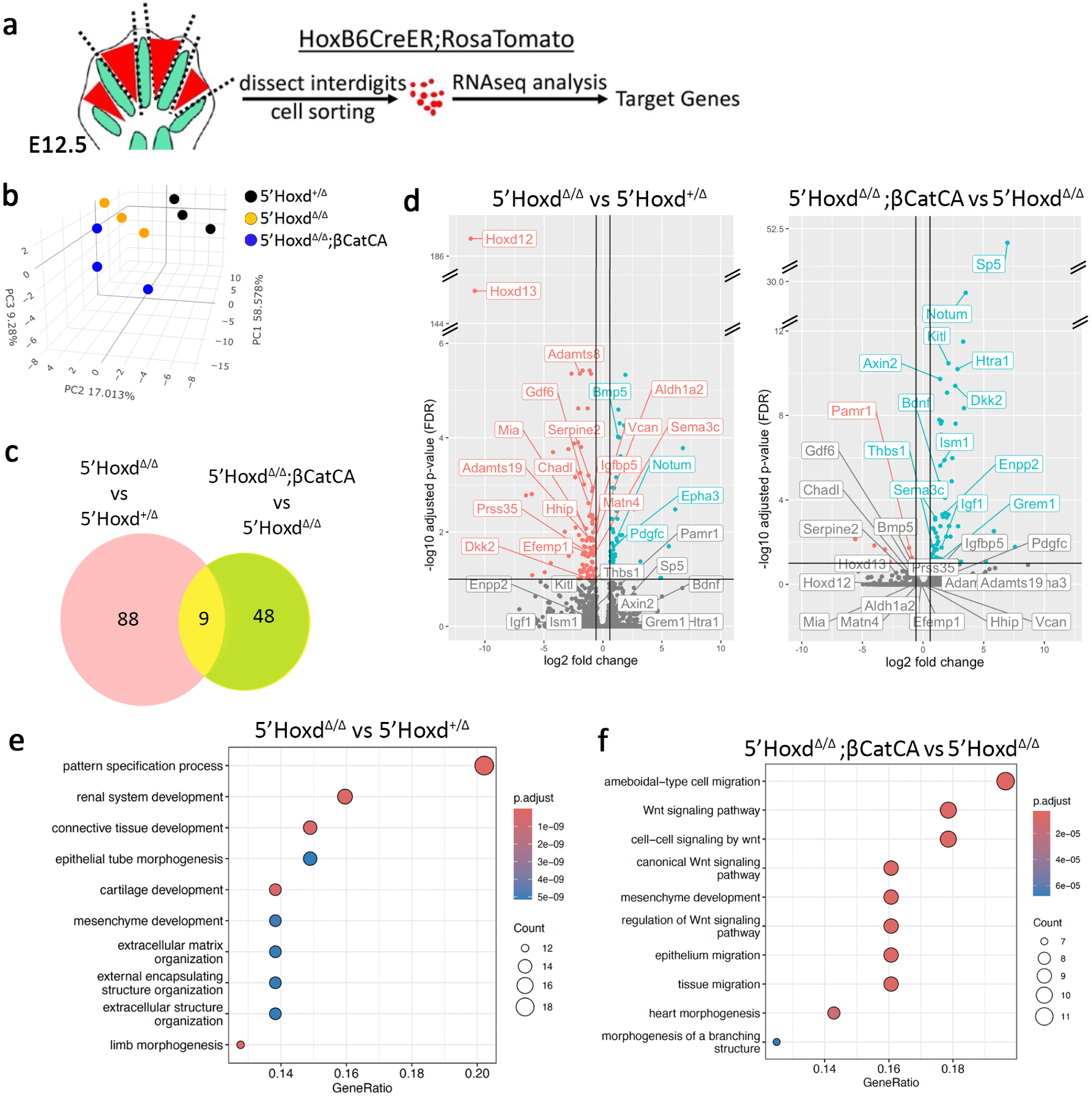
Transcriptome analysis identifies Wnt antagonists as candidate genes for restoring digit joint formation in 5’Hoxd^Δ/Δ^;βCatCA. **(a)** Experimental design of interdigit tissue isolation for transcriptome analysis. Tamoxifen was injected at Ell.25. **(b)** PCA analysis shows distinct segregation of each genotype which has three individual sample sets. **(c)** Venn diagram shows nine among 145 significant DE genes are overlapped between the 5’Hoxd^Δ/Δ^ vs 5’Hoxd^+/Δ^ and 5’Hoxd^Δ/Δ^;βCatCAvs 5’Hoxd^Δ/Δ^ comparison (FC>1.5; FDR<0.1). **(d)** Volcano plot visualization of extracellular genes in d are highlighted in positive DE (green), negative DE (red) and non-significant (gray) in the 5’Hoxd^Δ/Δ^ vs 5’Hoxd^+/Δ^ or 5’Hoxd^Δ/Δ^;βCatCAvs 5’Hoxd^Δ/Δ^ comparison (FC>1.5; FDR<0.1). **(e)** Gene-set GO biological process enrichment for top 97 DE genes in 5’Hoxd^Δ/Δ^ vs 5’Hoxd^+/Δ^ comparison. Gene-set GO biological process enrichment for top 57 DE genes in 5’Hoxd^Δ/Δ^;βCatCAvs 5’Hoxd^Δ/Δ^ comparison.

Comparisons between 5’Hoxd^Δ/Δ^ with 5’Hoxd^+/Δ^ and 5’Hoxd^Δ/Δ^;βCatCA with 5’Hoxd^Δ/Δ^ identified a total of 145 DE genes (FC ≥1.5; Padj (FDR) ≤ 0.1 filters), including 97 in 5’Hoxd^Δ/Δ^ vs 5’Hoxd^+/Δ^ control, 57 in 5’Hoxd^Δ/Δ^;βCatCA vs 5’Hoxd^Δ/Δ^, and 9 genes that were DE in both comparisons (Figure 3c, Table S1). Consistent with several previous genetic and profiling studies^12,30,31^, in the 5’Hoxd^Δ/Δ^ mutant compared to 5’Hoxd^+/Δ^ controls (Tables S1, S2), Bmp pathway components were upregulated (eg. *Msx2*; 1.75-fold, FDR=0.11 and *Bmp5*; 2.4-fold, FDR<0.01), *Hoxa13* displayed compensatory upregulation in the absence of *Hoxd13*, and several DE genes identified previously in microarray analysis of E12.5 5’Hoxd^Δ/Δ^ distal autopod behaved comparably in our analysis (*Papss2*, *Aldh1a2*, *Nr2f1* down-regulated; *Epha3, Tenm4/odz4* up-regulated).

Genes that were DE in both comparisons were of particular interest, considering that any factor in this group that normally plays a role in interzone/joint formation should be oppositely regulated in the 5’Hoxd^Δ/Δ^ mutant with or without rescue. Notably, this small set of 9 genes (Table S1, yellow rows) included only three expected to act non-autonomously, *Dkk2*, *Notum* and *Sema3c* (Table 1, yellow boxed genes in Table S1). *Sema3c* acts as either a chemorepulsive or chemoattractive agent to guide neural cell migration^32^, especially neural crest, but not in digit patterning^33^. The Wnt pathway is known to play key roles ranging from early to late events including limb initiation/AER induction, through digit patterning and tissue morphogenesis ^4,7,17,34–37^. As expected, the 5’Hoxd^Δ/Δ^; βCatCA vs 5’Hoxd^Δ/Δ^ DE set included multiple known canonical Wnt targets (Figure 3d,f), such as *Sp5*, *Axin2*, and Wnt-antagonists involved in negative-feedback regulation (*Notum, Nkd1, Dkk2*). Of these, *Dkk2* reduction in both 5’Hoxd^Δ/Δ^ and elevation in βCatCA-rescued 5’Hoxd^Δ/Δ^ interdigits were highly significant. *Notum*, which depalmitoylates Wnt ligand extracellularly, was modestly upregulated in 5’Hoxd^Δ/Δ^ mutant, and strongly upregulated by βCatCA rescue (Table 1).

**Table 1.** Secreted factors differentially expressed in βCatCA rescue.

| Gene: | *FC in 5'Hoxd$\Delta/\Delta$<br>vs 5'Hoxd$^{+/\Delta}$: | FDR: | *FC in 5'Hoxd$\Delta/\Delta$; βCatCA<br>vs 5'Hoxd$\Delta/\Delta$: | FDR: |
| --- | --- | --- | --- | --- |
| Dkk2 | 2.3 | 0.09 | 6.3 | 4e-10 |
| Notum | 2.2 | 0.01 | 11.6 | 2.4e-30 |
| Sema3c | 1.7 | 0.03 | 2.0 | 6.9e-4 |
| Bdnf | NS<sup>†</sup> | NS | 2.5 | 1.4e-5 |
| Enpp2 | 1.9 | NS | 3.5 | 4.2e-4 |
| Grem1 | 1.2 | NS | 2.3 | 0.09 |
| Htr4a1 | 1.7 | NS | 7.0 | 6.4e-11 |
| Igf1 | 1.5 | NS | 2.8 | 0.02 |
| Ism1 | NS | NS | 2.6 | 2.4e-6 |
| Kitl | 1.7 | NS | 4.3 | 3.4e-11 |
| Pamr1 | 1.6 | NS | 1.9 | 0.06 |
| Thbs1 | 2.0 | NS | 2.1 | 0.08 |

\*red denotes fold-change reduced, blue denotes fold-change increased
<sup>†</sup>NS, non-significant

To comprehensively consider all factors that could act non-autonomously from interdigits to restore interzone/joint formation, all DE genes in the 5’Hoxd^Δ/Δ^;βCatCA vs 5’Hoxd^Δ/Δ^ comparison alone were filtered for extracellular location and function using GO ^38^ and David (https://david.ncifcrf.gov/tools.jsp) (but excluding structural collagen genes). In addition to the 3 genes that were DE in both comparisons, 9 DE genes acting non-autonomously as secreted signaling factors, antagonists, or enzymes were identified (Table 1, green boxed genes in Table S1). Some of these also displayed altered expression in 5’Hoxd^Δ/Δ^ vs control, but did not reach statistical significance. Most notable in this group was the Bmp antagonist *Grem1*, induced 2.4-fold by βCatCA. Several others are factors more involved in late morphogenesis (eg. vascular, neural regulators). HCR fluorescent in situ of the major candidate signaling factors identified (*Grem1, Dkk2, Notum*), as well as *Bmp5*, which is elevated in the 5’Hoxd^Δ/Δ^, and *Axin2*, a major canonical Wnt target induced by βCatCA rescue, confirmed the expression changes in mutant and in rescued mutant E12.5 limb buds predicted by the DE analyses (Table 1, Figures 3d, S3).

### Genetic manipulation of Wnt signaling alters Bmp activity levels and digit joint formation

Because both Bmp and Wnt pathways have been implicated in joint formation, we assessed whether the identified candidates in these pathways (altered by interdigit βCatCA expression) played a causative role in restoring joints in 5’Hoxd^Δ/Δ^ digits. The Bmp antagonist *Grem1* was increased by βCatCA (Table 1) and would be expected to inhibit activity of multiple Bmp ligands. We tested if raising the interdigit *Grem1* level could restore digit joint formation in 5’Hoxd^Δ/Δ^ digits by activating *RosaGrem1* with Hoxb6CreER ^39^. Consistent with previous results ^12^, although increasing *Grem1* expression alone in 5’Hoxd^Δ/Δ^ had a modest effect on phalanx length, it did not improve digit joint formation in the mutant (n=12/12, Figure S4). Expression of canonical Wnt targets *Notum* and *Dkk2* were increased by interdigital βCatCA. Both *Dkk2* and *Notum* knockout mice are viable and their limbs appear normal at birth^40,41^. However, both multiple Wnt ligands and Wnt antagonists (including *Dkk2*) are expressed in interdigits and might act redundantly^35,42^. Since *Dkk2* is selectively expressed in interdigits normally and was down-regulated in 5’Hoxd^Δ/Δ^ as well as being up-regulated by βCatCA rescue (Table 1, Figure 3d, S3), we tested if *Dkk2* removal could block the joint restoration by interdigit βCatCA in 5’Hoxd^Δ/Δ^ digits. Deletion of *Dkk2* in sibling controls did not alter digit morphology including joint formation (Figure 4e). Additionally, *Dkk2* deletion did not worsen impaired joint formation in 5’Hoxd^Δ/Δ^ mutants (n=4/4, Figure 4a,d). However, *Dkk2* removal did prevent joint restoration by βCatCA in 5’Hoxd^Δ/Δ^;βCatCA embryos (n=19/34, Figure 4b,c-brackets). Together, these results suggest that interdigit βCatCA rescues digit joint formation in the 5’Hoxd^Δ/Δ^ by inducing negative feedback targets of the Wnt pathway, including secreted Wnt antagonists, such as *Dkk2*. This implies that reducing, rather than activating, Wnt activity in progenitor cells contributing to the PFR plays an early role in joint specification.

**Figure 4.**
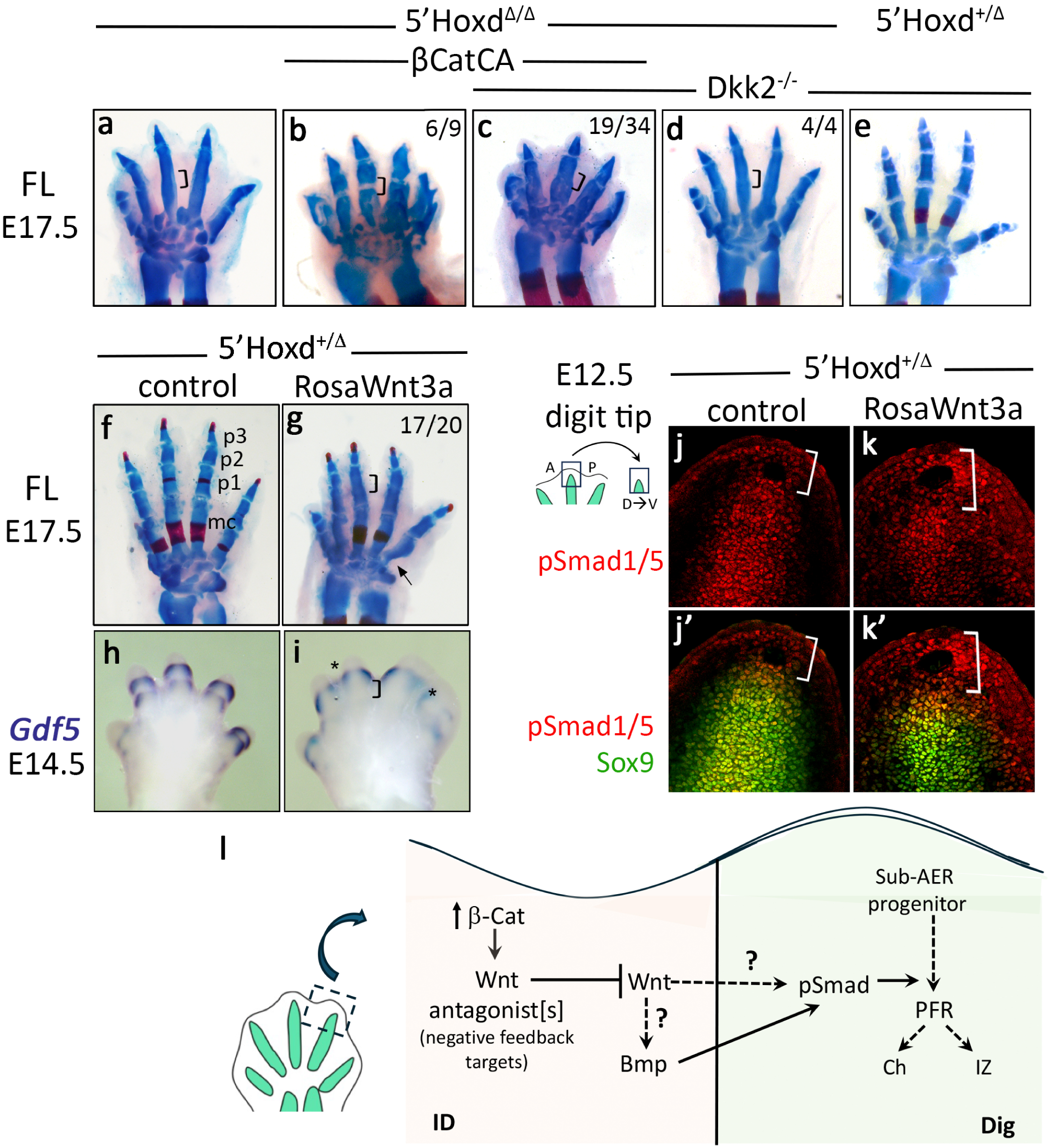
Effects of elevating Wnt signaling on digit joint formation in 5’Hoxd^+/Δ^ control and in βCatCA-rescued 5’Hoxd^Δ/Δ^. **(a-e)** E17.5 skeletal stain showing effect of Dkk2 loss on digit joint formation in βCatCA-rescued 5’Hoxd^Δ/Δ^ (compare b and c) and in control (e). Tamoxifen injected at Ell.25 for Hoxb6CreER activation of βCatCA expression. Bracket indicates presumptive (a,c,d) or rescued (b) digit joint region. Numbers in panels indicate the frequency of phenotype shown in the limbs analyzed. **(f-i)** Hoxb6CreER-activated Wnt3a expression in 5’Hoxd^+/Δ^. Tamoxifen injected at E10.75. **(f, g)** skeletal stain of E17.5 digits, **(h,i)** *Gdf5* in situ hybridization to visualize E14.5 interzones. Bracket shows loss of pl/p2 digit joint, *indicates syndactyly. Arrow: indicates altered digit morphology. **(j-k)** pSmadl/5 immunostaining of E12.5 distal digit tip section (dorsoventral, as shown in schematic). Brackets indicate sub-AER mesenchymal region. Sox9 immunostaining visualizes chondrogenic regions. **(I)** schematic summary of possible Wnt ligand and Wnt antagonist effects on Bmps, pSmad activity, acting downstream of interdigit βCatCA. See text for details, me, metacarpal bone; pl-p3, phalanges 1-3.

To further test this idea, we checked whether elevating Wnt levels modulates digit joint formation. Activating a conditional *RosaWnt3a* transgene using Hoxb6CreER and early tamoxifen injection at E10.75 to achieve broad, robust expression caused frequent digit joint loss in 5’Hoxd^+/β^ embryos (n=17/20, Figure 4g), and rarely also in wild-type (5’Hoxd^+/+^) embryos (n=1/10). Enforced Wnt ligand expression also caused mild syndactyly and altered cartilage morphology (Figure 4g,i), and *Gdf5* expression was absent in presumptive interzone joint regions during early-stage joint development (Figure 4h,i brackets). These results raised the possibility that high canonical Wnt activity in cells transiting into the PFR may act to suppress interzone specification, similarly to Bmps. Indeed, pSmad1/5 levels were correspondingly increased in the sub-AER mesenchyme and extending into the PFR region in Wnt3a transgenic embryos compared to sibling controls (Figure 4j,k brackets). These results indicate that Wnt signaling cooperates with the Bmp pathway to elevate pSmad1/5 activity in progenitor cells transiting to PFR and thereby promote chondrogenic at the expense of interzone specification, suggesting that interdigit Wnt antagonists act to maintain interzone/joint specification by modulating digit tip Wnt signaling (Figure 4l).

### Wnt signaling enhances Bmp activity by destabilizing Gsk3

Cooperation between Bmp and Wnt pathways has been previously observed in some contexts, for example, ventralization of mesoderm during Xenopus gastrulation, where it was previously shown that canonical Wnts synergize with Bmp activity by stabilizing Bmp receptor-activated Smads. Gsk3β, acting together with MAPK, destabilizes Smad1 protein by phosphorylating the Smad linker region, which targets it for ubiquitination and degradation^43,44^(see Figure 5). By inactivating Gsk3β, canonical Wnts can stabilize pSmad1 and enhance Bmp activity, raising the possibility that interdigit Wnt antagonists act downstream of βCatCA to stabilize Gsk3β and reduce Bmp response by promoting pSmad1/5 linker phosphorylation and turnover (Figure 5e). To test this hypothesis, we used pharmacologic approaches in short-term limb bud organ culture (Figure 5a) to block either Gsk3β, the ubiquitin ligase Smurf1, or proteasome activities, and assess whether the clearance mechanism for Smad1 protein in Xenopus is conserved in the mammalian digit-forming region. We assayed Bmp response in short-term cultures of E12.5 limb buds by measuring both pSmad1/5 protein (C-terminal activating phospho-serine) and the direct Bmp target *Msx2*. Short-term culture did not, by itself, alter *Msx2* expression compared to non-cultured, contralateral limb buds (Figure S5). Treatment with either a Gsk3 inhibitor (CHIR 99021 or Bio), Smurf1 inhibitor (A01), or proteasome inhibitor (MG132), all increased *Msx2* expression over vehicle-treated controls (Figures 5c, S5). Likewise, elevated pSmad1/5 protein was observed in the distal digit tip of CHIR 99021 treated limb buds (Figure 5d). In contrast, treatment with Wnt-C59, an inhibitor of porcupine and Wnt ligand secretion, resulted in reduced pSmad1/5 protein and *Msx2* RNA levels compared to controls (Figure 5d). These results suggest that Wnt signaling enhances Bmp activity in limb bud digit tips by modulating Gsk3β to stabilize pSmad1/5 proteins (Figure 5e).

**Figure 5.**
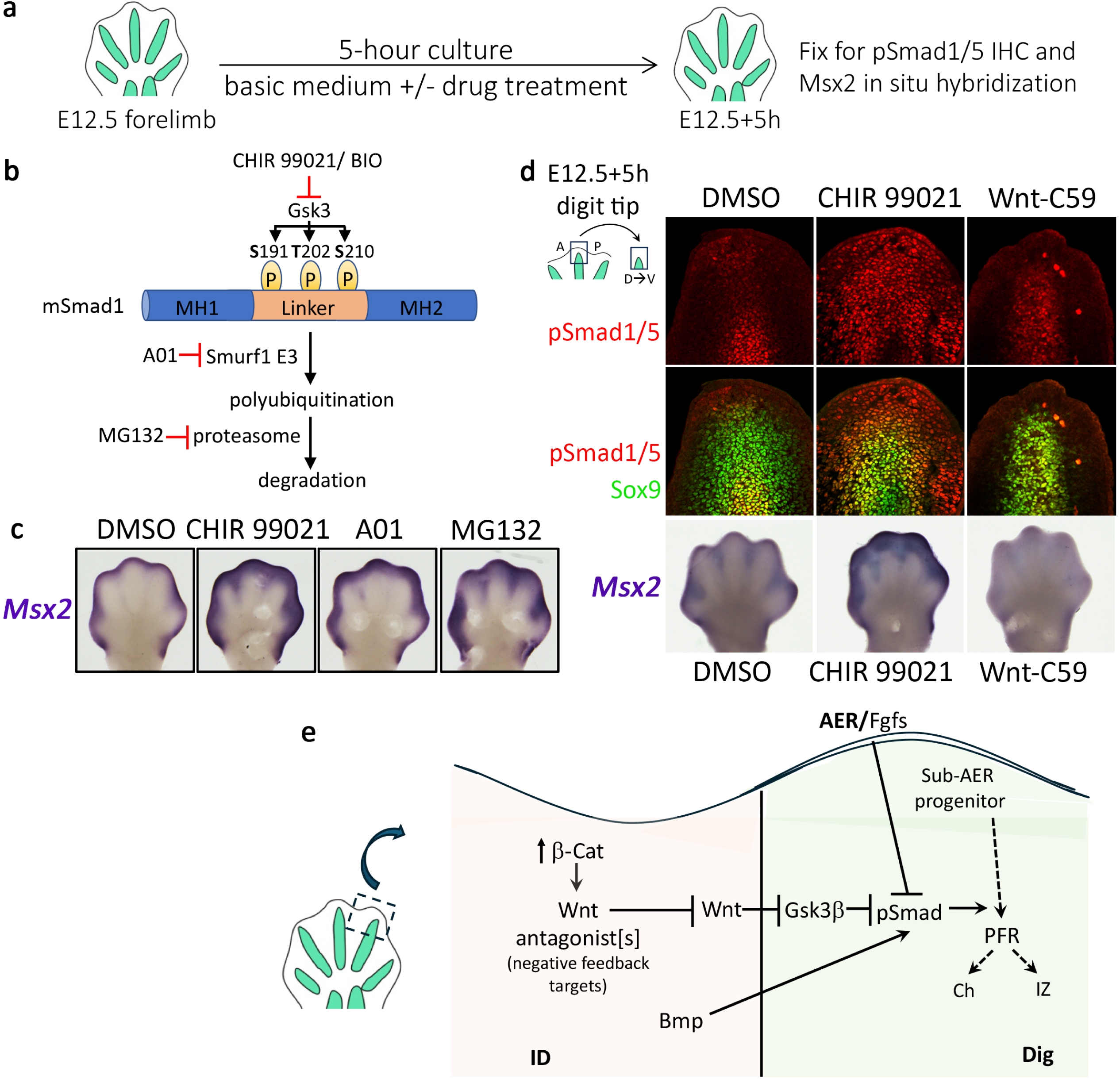
Wnt signaling enhances distal Bmp activity by reducing Gsk3. **(a)** Experimental design of short-term E12.5 limb bud organ culture, **(b)** Schematic showing site of action of pharmacological agents used. Modified from ref. (43). **(c)** in situ hybridization of direct Bmp target, *Msx2*, shows enhanced Bmp activity after inhibition of Gsk3 activity, or inhibition of proteasome activity, **(d)** Upper panels show pSmadl/5 and Sox9 staining in digit tips at El2.5+5h (dorsoventral section, as shown in schematic). Inhibiting Gsk3 activity increases accumulation of pSmadl/5 in distal digit tip and conversely, inhibiting Wnt ligand secretion reduces number of pSmadl/5-positive cells. Bottom panel shows changes in Bmp response visualized by whole mount *Msx2* expression, **(e)** schematic summary of effect of Wnt pathway inhibition (pharmacologic invitro, or by Wnt antagonists in vivo) on Gsk3 activity. Gsk synergizes with AER-Fgfs to destabilize pSmads through linker phosphorylation (b). Wnts synergize with Bmp pSmad-activation by promoting Gsk3 degradation. See text for details.

## Discussion

Previous work indicates that Wnts prevent mesenchymal cells from adopting a chondrogenic fate by inducing stable epigenetic silencing of the *Sox9* locus via DNA-methylation. However, Fgfs act to prevent this and maintain the *Sox9* promoter in a poised state^6^. In this study, we discovered that Wnts can augment Bmp activity by suppressing Gsk3β to enhance pSmad1/5 stabilization, resulting in mesenchymal cells entering a pre-chondrogenic state precociously. This unexpected Wnt-Bmp synergy is held in check by Wnt antagonists, suggesting that Wnts regulate pre-chondrogenic fate specification at several levels with opposing effects. An appropriate balance between Wnt and Wnt-antagonist activities maintains the *Sox9* locus primed for expression while also enabling early pSmad1/5 accumulation in response to Bmps to promote pre-chondrogenic mesenchymal cell compaction. Wnt antagonists play key roles both in preventing precocious or excessive accumulation of pSmad1/5 by stabilizing Gsk3β in cells when Fgf levels are high, and in preventing *Sox9* silencing as Fgfs decline by curtailing excess Wnt activity (Figure 6).

**Figure 6.**
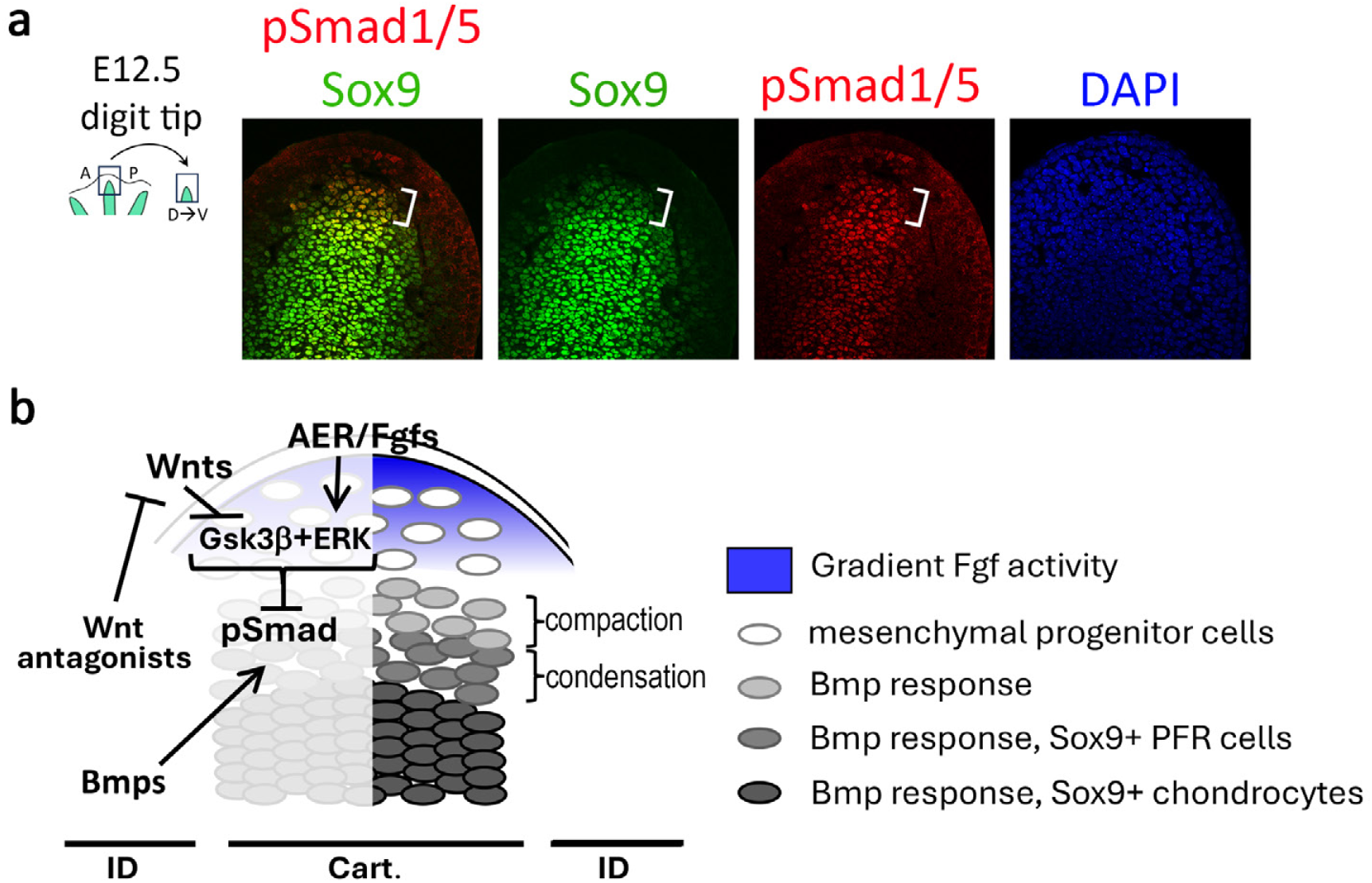
Bmp activation precedes Sox9 expression as mesenchymal cells transit into PFR cells. **(a)** Sox9 and pSmadl/5 staining in wild-type E12.5 limb bud distal digit tip sections (dorsoventral, as shown in schematic). The bracket indicates condensing pre-PFR region where Bmp activation of pSmadl/5 initiates, followed by low level Sox9 expression. **(b)** Interaction of signaling pathways regulating the progression of mesenchymal progenitor cells to chondrocytes. See discussion for details.

The two major phenotypes present in the 5’Hoxd^Δ/Δ^, digit shortening with phalanx loss (brachydactyly) and digit joint loss, result from a perturbed balance between the Fgf, Wnt and Bmp pathways. Digit interzone specification is initiated in PFR cells when the Bmp response is reduced ^12^. Here we show that interdigit-activated βCatCA rescues joint formation non-autonomously in 5’Hoxd^Δ/Δ^ digits by inducing expression of canonical Wnt targets that act as negative feedback regulators, including *Dkk2,* which is down-regulated in the mutant. This suggests that digit joint loss in the 5’Hoxd^Δ/Δ^ results, at least in part, from reduced Wnt antagonist levels and consequent elevated Wnt activity. This impression is supported by results of *Wnt3a* overexpression in wild-type embryos during phalanx formation, which also results in excess pSmad1/5 accumulation and digit joint loss. These results all imply that high Wnt activity impedes a fate switch to interzone specification. However, previous work has led to the proposal that canonical Wnt/βCatenin signaling promotes interzone formation^4,17^. Although overexpressed Wnts induced ectopic interzone formation with *Gdf5* expression, *βCatenin* deletion, or selective mesenchymal deletion of *c-Jun*, a transcriptional activator of Wnts in interzone ^17,20,45^, did not abolish initial *Gdf5* expression in the emerging interzone of mutant mice. *Gdf5* expression only declined later, giving rise to joint fusion phenotypes. The loss-of-function results indicate that interzones are still specified in these mutants and that canonical Wnt/βCatenin signaling is critical for subsequent interzone maintenance and maturation into joints, rather than initiation. Notably, our previous study indicated that down-regulation of Bmp activity in 5’Hoxd^Δ/Δ^ can restore both phalanx and joint formation. However, interdigit βCatCA restores only joint formation but not phalanx loss in 5’Hoxd^Δ/Δ^ digits. Further work will be required to determine what additional factors regulating digit mesenchymal progenitor maintenance or expansion may be altered following Bmp pathway down-modulation to restore phalanx formation in 5’Hoxd^Δ/Δ^ digits.

In the case of interzone restoration by interdigit βCatCA, induced Wnt antagonists counteract synergistic effects of canonical Wnt activity on pSmad stimulation by Bmps in sub-AER progenitors, resulting in biphalangeal digits with rescued joints. By removing Gsk3β activity, elevated Wnts enhance pSmad1/5 accumulation and thereby accelerate the pace of transition from mesenchymal progenitors to PFR cells and chondrogenic commitment, consequently impeding the fate switch to interzone formation as seen in 5’Hoxd^Δ/Δ^, or following Wnt3a misexpression, and leading to joint loss. Conversely, reducing either Bmp activity ^12^ or Wnt activity in 5’Hoxd^Δ/Δ^ can restore digit interzone/joint formation.

Wnts and Bmps have been proposed to play key roles in promoting digit regeneration^46,47^, and it is important to discern how blastema cells, the presumptive regenerative progenitor cells, integrate pathway activities to achieve full regeneration of both morphology and function. In this study, using genetic approaches, we uncovered crosstalk between Bmps and Wnts and discovered that Wnt antagonists play a pivotal role in maintaining progenitor plasticity during the transition to PFR and phalanx formation in embryonic digits. These results shed light on how complex signaling interactions are balanced to regulate different specific digit identities (morphologies and phalanx/joint numbers) and help advance our understanding of how complex signaling interactions modulate cell fate determination during morphogenesis.

## Acknowledgements

We thank Marian Ros for critical comments on the manuscript, Ryan Kelly for assistance with GO analysis, and Denny Duboule/Jozsef Zakany, Gail Martin, Mark Taketo, Steve Vokes, and Terry Yamaguchi for generously providing mouse lines.

## Author Contributions

BL and SM designed the project and wrote and edited the MS, BL performed experiments, EK and MP performed late stage analysis of skeletal joint formation and MS editing, and SD analyzed all the RNAseq data.

## Methods

### Mouse Strains and Embryo analyses

All animal studies were carried out according to the ethical guidelines of the Institutional Animal Care and Use Committee (IACUC) at NCI-Frederick under protocol #ASP-23-405. The *Catnb^+/^*^C-exon3 21^, *Hoxd^Del^*^(11–13)^*^/+^* (*5’Hoxd*^+/β^) ^23^, *NogginLacZ* ^2^, *RosaGrem1* ^39^, Hoxb6CreER ^26^, OsrCre ^28^, *Dkk2^−/−^*(JAX #030130) ^41^, *Sox9CreER* ^48^, *RosaWnt3a* ^49^, and *Rosa-tdTomato* ^50^ alleles used have all been reported previously. For timed matings, noon on the day of post-coital plug was considered to be E0.5. For inducible Cre-drivers, a single dose of 3mg tamoxifen (in all cases) was injected intraperitoneally as previously described ^51^ at the time indicated.

### Skeletal preparation

Embryos were collected and fixed in ethanol, followed by acetone dehydration. The skeletons were stained in 0.3% Alcian Blue 8GS (Sigma-Alderich #A5268) and 0.1% Alizarin Red S (Sigma-Alderich #A5533) in 70% ethanol containing 5% acetic acid. Stained tissues were cleared in 2% potassium hydroxide and transferred to 50% glycerol for imaging ^12^.

### Sectioned in situ hybridization analysis

This procedure was carried out as described previously ^52^. Briefly, limbs were embedded in paraffin, and sectioned. Serial 5μm-thick sections were pretreated with 1μg/ml proteinase K (Sigma-Aldrich #SAE0009) in 50mM Tris-HCl, 5mM EDTA pH 7.5 for 1 min at room temperature, immediately post-fixed in 4% paraformaldehyde buffer for 10 min, followed by washes and treatment with 0.25% acetic anhydride in triethanolamine buffer for 15 min. Sections were hybridized with antisense or sense [^35^S]-labeled riboprobes (approximately 1 x 10^6^ DPM/section) at 50°C for 16 h. After hybridization, slides were treated with 20 μg/ml RNaseA for 30 min at 37°C and dehydrated in ethanol, then coated with Kodak NTB-3 emulsion diluted 1:1 with water, air-dried and exposed in a light-proof box for image development. cDNA clones used for two kinds of mouse Collagen Type II are: a 121 bp collagen IIA (284–404, NM_031163); a 356 bp collagen IIB (2409–2764, NM_031163)^53^.

### Whole mount colorimetric and Hybridization chain reaction (HCR) in situ analysis

Embryos were collected, fixed in 4% paraformaldehyde, and stored in 100% methanol until further analysis. The fixed embryos were bleached in 5% hydrogen peroxide in methanol followed by rehydration and a brief proteinase K treatment (5-15 min., 20ug/ml). For colorimetric in situ hybridization, embryos were hybridized with digoxigenin-UTP-labeled antisense riboprobes in a standard hybridization buffer containing 50% formamide, 1%SDS, 0.75M NaCl at 70°C overnight. Embryos were washed in hybridization buffer at 70°C, hybridization buffer with reduced salt (0.15M NaCl) at 50°C, and in reduced salt (50mM NaCl) without formamide at 70°C, and then transferred to TBST (25mM Tris pH 7.4, 150mM NaCl, 0.1% Tween20) for incubation with anti-digoxigenin antibody conjugated to alkaline phosphatase (1:2000, Roche #11093274910) at 4°C overnight. The colorimetric reaction was developed in BM Purple (Roche #11442074001) at room temperature until a clear signal was present. The whole mount HCR in situ hybridization followed the recommended procedures for third generation HCR probes ^54^ designed by Molecular Instruments (Los Angeles, CA) and HCR staining was analyzed by confocal microscopy.

### Immunofluorescence imaging

Embryos were collected, fixed in 4% paraformaldehyde and stored in 70% ethanol with phosphatase inhibitor cocktail (Millipore #524631) until use. Forelimb samples for tdTomato labeled lineage tracing analysis were embedded in OCT, followed by cryo-sectioning at 10um thickness. Cryo-sections were stained with Alexa Fluor 488 conjugated anti-Sox9 antibody (1:500, Abcam #ab196450). For other immunofluorescent staining, fixed limb buds were embedded in 7% low-melting agarose for vibratome sections. 100um sections were treated with anti-phosphoSmad1,5 (1:200, Cell Signaling #9516) and visualized with Alexa Fluor 594 secondary antibody, followed by counterstaining with Alexa Fluor 488 conjugated anti-Sox9 antibody to visualize chondrogenic condensations. Stained samples were imaged by confocal microscopy. At least 3 independent samples were imaged for analysis in each genotype.

### Short-term limb bud culture with pharmacological agents

Embryonic stage E12.5 mouse forelimb buds were dissected in iced cold PBS, followed by incubation in serum-free limb bud minimal media ^55^ either with DMSO vehicle alone (0.01%, Sigma-Aldrich #8418), CHIR 99021 (10uM, Tocris #4423), BIO (5uM, Tocris #3194), A01 (5uM, MedChemExpress #HY-110195), MG132 (50uM, Tocris #1748) or Wnt-C59 (10uM, Tocris #5148) at 37C, 5% CO2 condition for 5 hours. After 5 hours, cultured limb buds were fixed in 4% paraformaldehyde and stored in 100% methanol for subsequent analysis by immunofluorescence staining and HCR as described above. At least 5 independent limb buds were treated and analyzed for each condition.

### Bulk RNAseq analysis

To generate different genotypes for transcriptome comparison, Hoxb6CreER^tg/tg^;5’Hoxd^+/Δ^ males were crossed with Catnb^+/C-exon3^/5’Hoxd^Δ/Δ^/Rosa-tdTomato^tg/tg^ females. Pregnant females were injected with Tamoxifen at E11.25 and embryos were collected at early E12.5 and held at 4C in PBS during rapid PCR genotyping. For each genotype of sibling embryos (5’Hoxd^+/β^, 5’Hoxd^Δ/Δ^ or 5’Hoxd^Δ/Δ^;βCatCA), forelimb buds of at least 3-4 embryos from two litters were grouped. Interdigit tissues [between d2-d3, d3-d4, and d4-d5, see Fig. 2a] were dissected with tungsten needles and dissociated with 0.25% Trypsin (Gibco #15090046). After 3-4 minutes incubation at 37C followed by gentle trituration, trypsin was inactivated with 1mM AEBSF (MP Biomedicals #193503) followed by addition of 1% serum and isolation of tdTomato labeled interdigit cells by FACS. Three independent experimental sets were collected and processed for bulk RNAseq. Barcoded libraries prepared from 9 samples of cDNAs were pooled and sequenced for paired-end 103bp reads using Illumina TruSeq, and unique reads were aligned to mouse GRC38/mm10 mm. Differential expression (DE) analysis was carried out using DESeq2 ^56^ on R version 4.3.2. The DAVID Functional Annotation Tool (https://david.ncifcrf.gov/tools.jsp) was used to identify secreted/extracellular factors in the DE gene sets and Gene Ontology for biological process enrichments analysis was carried out with Pathview package version 1.46.0 ^38^.

### Data Availability

All data are available from authors. All raw sequence data from bulk RNAseq experiments will be deposited in SRA/NCBI upon acceptance for publication.

---

Supplemental Information for excel data spreadsheets (Tables S1, S2):

**Table S1.** Normalized counts and DE analysis for all genes included in Figure 2C – Venn diagram summary (FC of >l.5x increase or decrease, AdjP(FDR)<0.1), and grouped/shaded to reflect Venn diagram colors: red rows: DE in 5’Hoxd^Δ/Δ^ vs 5’Hoxd^+/Δ^; green rows: DE in 5’Hoxd^Δ/Δ^;βCatCA vs 5’Hoxd^Δ/Δ^; yellow rows: DE in both comparisons. Boxed genes symbols indicate secreted factors in each of the comparison categories.

**Table S2.** Complete data set of normalized counts for all genes in which the maximum expression (max expr) counts in at least 1 of the 9 biological replicates reached a value > 10.

Shorthand headings for genotypes if interdigit samples analyzed in Supplementary Tables1,2 [for each genotype - 3 biological replicates analyzed]:

Del_Het = 5’Hoxd^+/Δ^

Del_Hom = 5’Hoxd^Δ/Δ^

Del_Hom_Exlox = 5’Hoxd^Δ/Δ^;βCatCA

Shorthad headings for DE analyses in Supplementary Tables1,2:

HomVsHet = 5’Hoxd^Δ/Δ^ vs 5’Hoxd^+/Δ^

bCatVsHom = 5’Hoxd^Δ/Δ^;βCatCA vs 5’Hoxd^Δ/Δ^

bCatVsHet = 5’Hoxd^Δ/Δ^;βCatCA vs 5’Hoxd^+/Δ^ [comparison only included in Suppl. Table 2]

**Figure S1.**
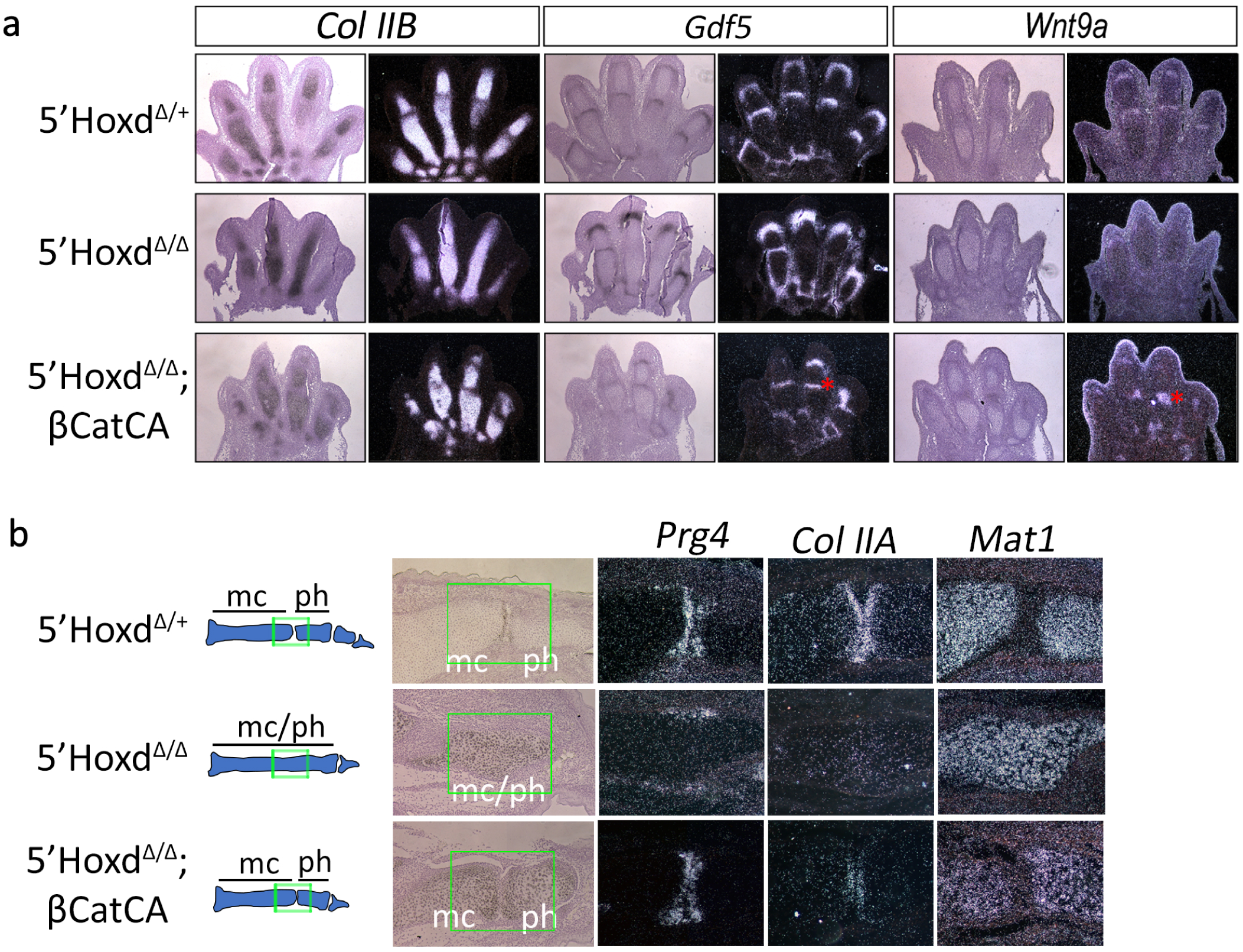
βCatCA expression in interdigits restores joint formation in 5’Hoxd^Δ/Δ^. **(a)** In 5’Hoxd^Δ/Δ^ mutant E14.5 digits, expression of *Gdf5* and *Wnt9a* in the interzone are absent, but are restored by activation of βCatCA in the interdigits using Hoxb6CreER (tamoxifen treatment at Ell.25). *Collagen type HB (Col HB)* is expressed in condensed cartilage^53^. * restored interzone. **(b)** In 5’Hoxd^Δ/Δ^ mutant PO (newborn) digits, *Proteoglycan 4 (Prg4)* is only expressed laterally at the periphery of the presumptive joint region, but is restored within the articular cartilage by activation of βCatCA in the interdigits using OsrCre, indicating normal joint maturation. *Col IIA,* which detects Exon 2, is expressed in articular cartilage^53^ but is absent in 5’Hoxd^Δ/Δ^ mutants, ph: phalanx; me: metacarpal bone.

**Figure S2.**
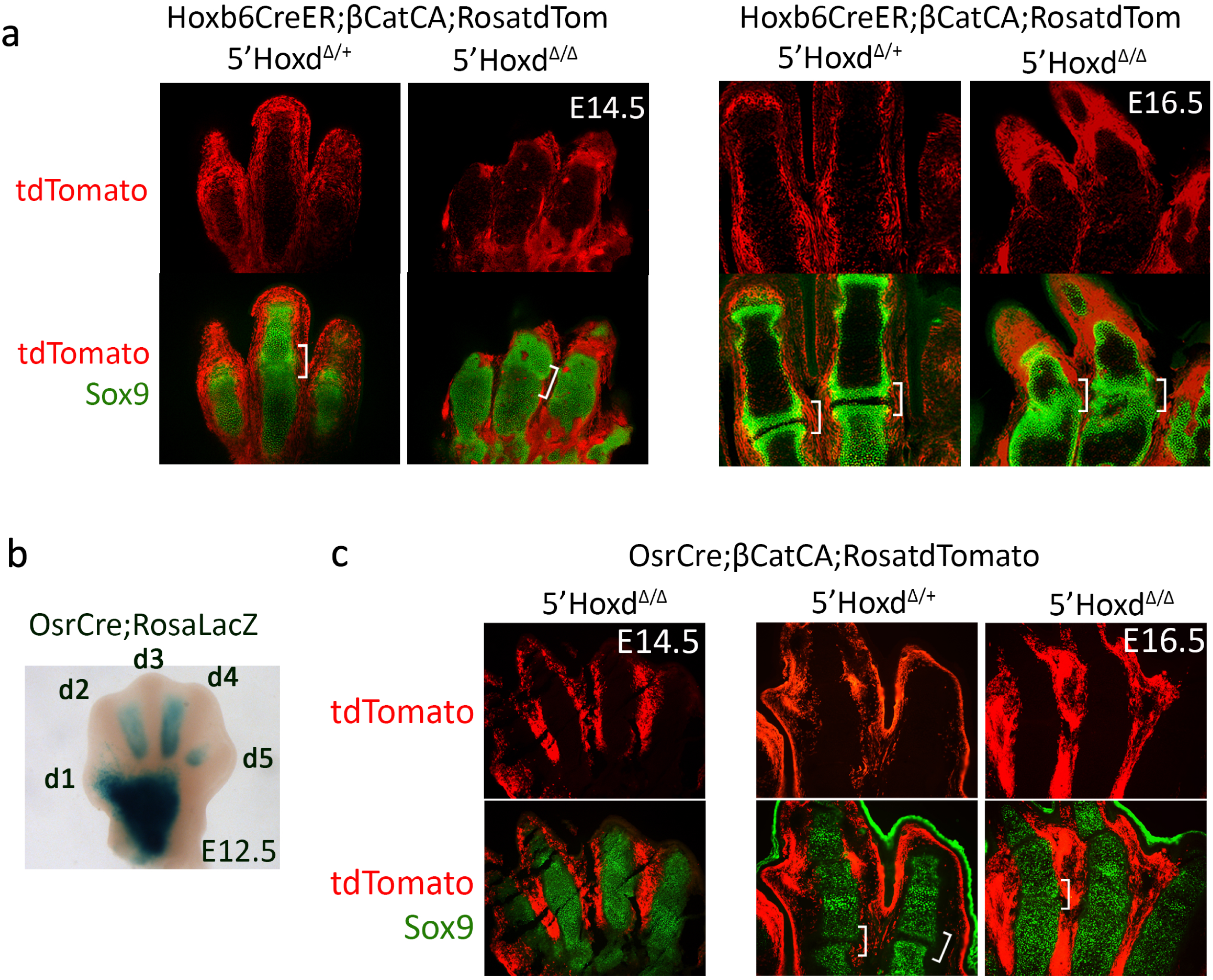
Lineage analysis shows that βCatCA expressing cells do not contribute significantly to cartilage or joint region after Hoxb6CreER activation and are absent from rescued digit3 joints in OsrCre-activated βCatCA. **(a)** Hoxb6CreER;Rosa-tdTomato labeled cells in 5’Hoxd^Δ/Δ^;βCatCA embryos are present predominantly in interdigit tissues with rare Tomato-expressing cells in cartilage observed at E14.5 or E16.5, following tamoxifen treatment at Ell.25.. Images are optical sections from 2OOum vibratome section stained in whole mount for Sox9 protein. **(b)** OsrCre activity is strongly expressed from the interdigits flanking digit 3 (d3) condensation by E12.5 in forelimb, but weakly expressed in other interdigits at this stage. **(c)** OsrCre;Rosa-tdTomato labeled cells in 5’Hoxd^Δ/Δ^;βCatCA embryos are observed only in interdigit tissues. Images from 5um cryo-sectons stained for Sox9 protein. Brackets indicates metacarpophalangeal joints. Note lack of contribution of Cre-activated βCatCA expressing cells to rescued interzone/joint.

**Figure S3.**
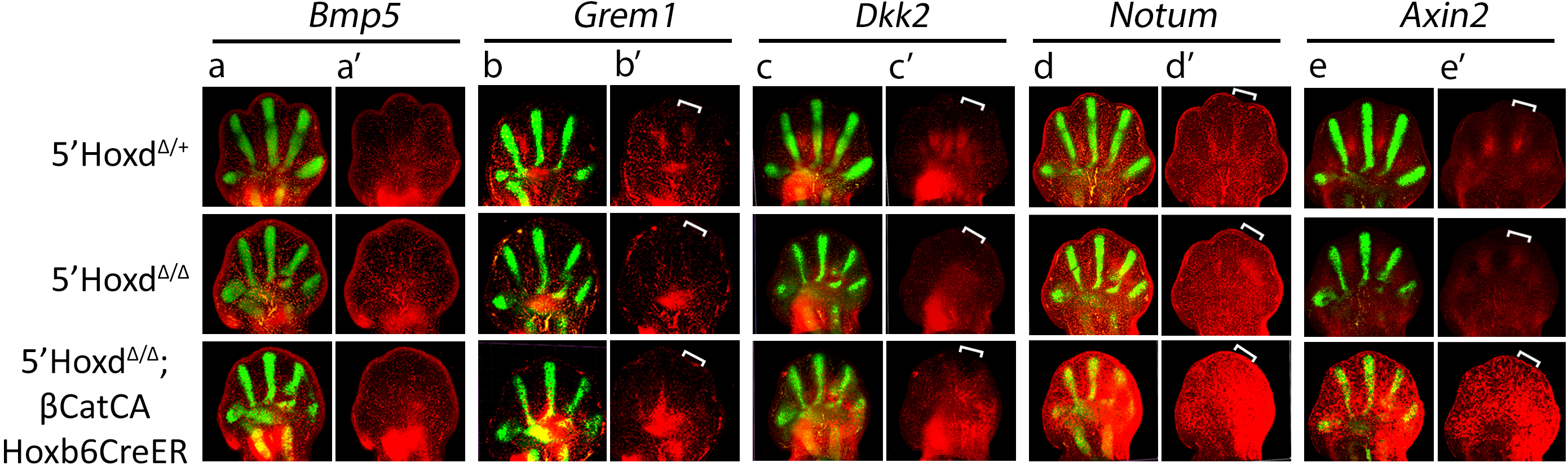
HCR in situ staining of Bmp and Wnt pathway DE RNAs identified in transcriptome analysis. **(a-e)** merged images of fluorescent HCR in situ staining for Sox9 and other genes as indicated. **(a’-e’)** fluorescent HCR in situ staining for DE genes alone. Brackets indicate interdigit tissues between digit 3-4 to highlight differences in DE gene expression between control, 5’Hoxd^Δ/Δ^ and βCatCA-rescued 5’Hoxd^Δ/Δ^ mutant.

**Figure S4.**
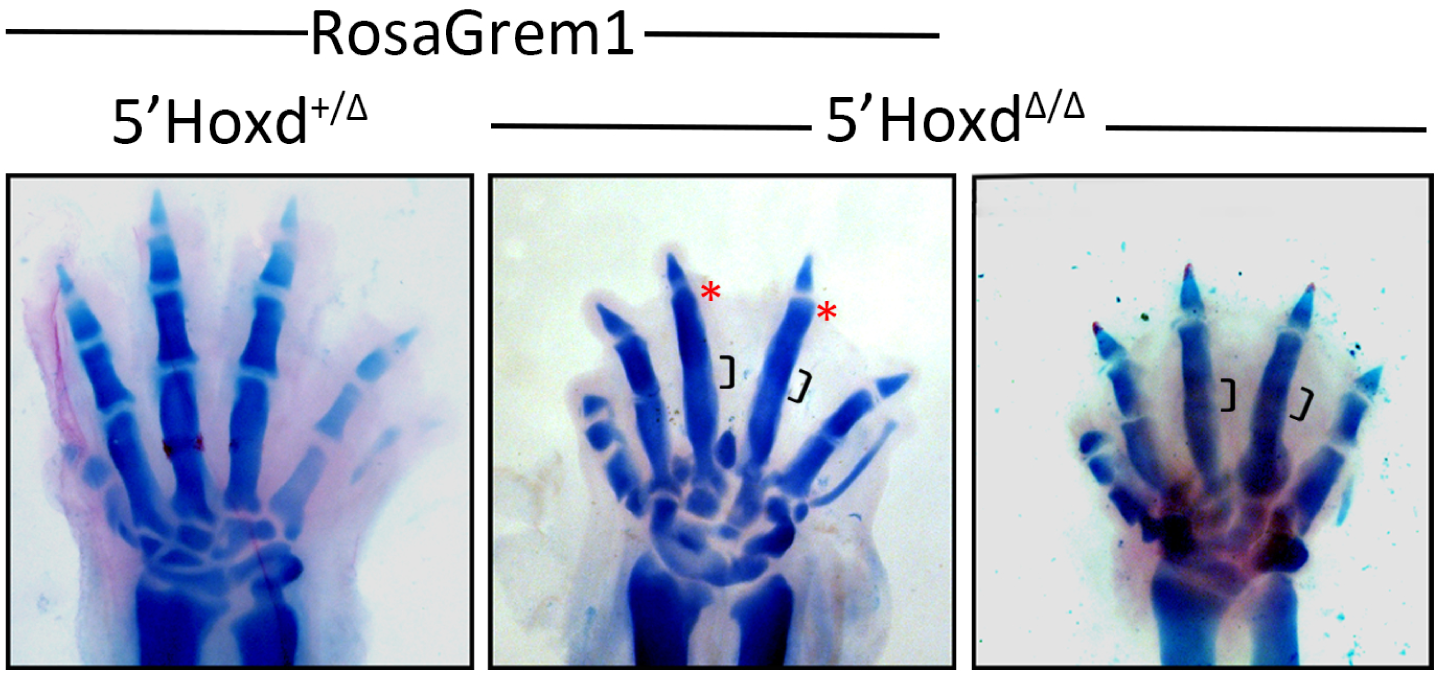
Transgenic ere-activated Grem1 misexpression in 5’Hoxd^Δ/Δ^ does not restore digit joint formation. Effect of RosaGrem1 activation with Hoxb6CreER on different 5’Hoxd genotypes as indicated. Tamoxifen injected at E10.75 and embryos collected at E16.5 for skeletal analysis (shown). * indicates modest effect on digit length. Brackets indicate presumptive joint region.

**Figure S5.**
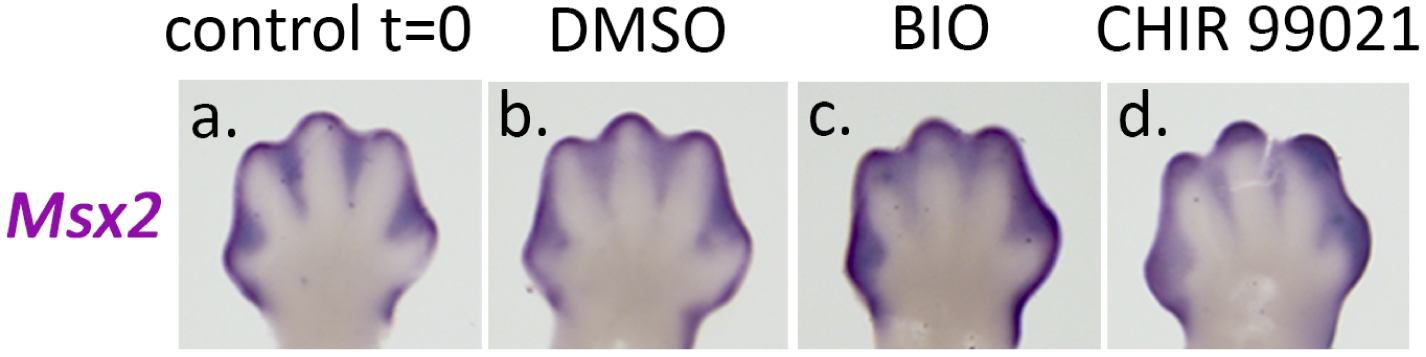
Bmp activity levels during extent of short-term limb bud organ culture. Bmp activity is not significantly altered during short-term limb bud organ culture (compare b with a) but is augmented by inhibition of Gsk3 activity during culture (compare c, d with b).

